# Impacts of Virtual Reality Experiences: Enhanced Undergraduate Student Performance and Engagement with use of 360-degree video

**DOI:** 10.1101/2023.02.16.528835

**Authors:** Erica Vola, Rebecca Stoltz, Charles A. Schumpert

## Abstract

Virtual reality has been used for a variety of training and gaming purposes. Recent studies have demonstrated their use in higher education enhances student engagement and positively impacts student performance. In this study, we implemented 360-degree video in an introductory biology lab and assessed student attitudes, engagement, and performance on various assessments as well as the overall course performance. Students were extremely positive about virtual reality, had minimal issues handling the technology, and indicated that engagement with the topic was better with virtual reality. Student performance increased on 2 of the 3 formative assessments analyzed, 1 of 1 summative assessment analyzed, and overall course grades in semesters with virtual reality were higher than in semesters without virtual reality. These results suggest virtual reality in higher education may enhance not only student engagement and performance, but also enhance student attitude about the topic.

## Introduction

Virtual reality (VR) is the computer-generated simulation of a three-dimensional image or environment that can be interacted with in a seemingly real or physical way. Over the past decade, use has rapidly evolved and has become more prevalent as technology advances. The first widely known experiments using head-mounted displays were conducted by Ivan Sutherland and showed participants different software based on the direction of the gaze (Sutherland, 1968). Now, VR has adapted with the fast-growing world to bring forth modern innovations including more portable forms of VR and augmented reality glasses that allow for interfacing of the digital and physical worlds. Studies have demonstrated a correlation between visual stimuli and related audio cues to enhance learning (Quak et al. 2015; Shams and Seitz, 2008). Utilizing VR in the classroom provides both visual and auditory cues to potentially provide enhanced value for learning (Huang and Liaw, 2018; Hu-Au and Lee, 2017).

Aside from the vast uses of VR in the entertainment and gaming industry, the development has proven efficient in the worlds of education and occupational training (Bennett and Saunders, 2019; Domingo and Bradley, 2018; Huang and Liaw, 2018; Hu-Au and Lee, 2017). The once novel concept of flight simulation is now standard in aviation training across the globe, with an increase in safety and a decrease in cost (Allerton, 2010). VR has also been implemented in military stress management training programs to decrease levels of perceived stress among soldiers (Pallavicini et al., 2016). Healthcare has played a key role in demonstrating the benefits of integrating VR with teaching and practicing. Orthopedic surgeons can improve their technical skills, and medical students can improve anatomy interest as well as test scores (Florence et al., 2016; Zhao et al., 2020). The benefits of VR in an educational setting stretch beyond performance and exam scores. University students using immersive versus desktop VR had significantly higher scores in presence and motivation which fall outside of strictly objective measurements (Liou and Chang, 2018; Makransky and Lilleholt, 2018).

Positive student outcomes rely heavily on not only the amount but also the quality of participation that is given in the classroom setting. Few concepts are as emphasized as student engagement in discussions surrounding how to best improve student test scores and classroom experiences (Axelson and Flick, 2010; McClenney, 2007; Mushtaq et al., 2018). Recent studies surrounding electronic learning demonstrate the direct correlation between test grades and the amount of interaction that a student has with the class (Mushtaq et al., 2018). As more studies demonstrate the importance of student engagement, it is necessary to investigate not only the quantity of engagement but also how the quality of engagement affects test scores (Zhao et al., 2020). Students learn in a variety of ways and the quality of learning can be a subjective measure. Education specialists debate as to how to universally measure this value, but there are several models becoming more accepted. Generally, educational practices that involve more senses produce longer lasting and higher quality working memories (Crissien et al., 2019). VR has great potential to increase the quality of student education and improve test score through multi-sensory immersive experience without leaving the classroom (Quak et al., 2015; Shams and Seitz, 2008). The sustainability of the VR experiences is greater than field trips and more equitable, as not all students have the resources for extensive excursions that some field trips require.

One learning theory proposes four stages known as the learning cycle where the use of multisensory learning has been beneficial in the initial phase. These stages are gathering, reflection, creation, and active testing (Quak et al. 2015; Shams and Seitz, 2008). The gathering phase of learning was focused on in this study using VR to increase the amount of stimulation through multisensory processing. Multisensory processing is the interaction of simultaneous signals from various sensory receptors (Quak et al., 2015; Shams and Seitz, 2008). These signals can be processed and interpreted into a signal event or combined in various ways to function as a single signal in a process known as multisensory integration (Quak et al., 2015; Shams and Seitz, 2008). The digital age has led to a significant increase in the ways that people consume multisensory data, especially in the case of adolescents and young adults who are accustomed to constant multisensory processing (Domingo and Bradley, 2018; Shams and Seitz, 2008; Spence, 2020; Starkey, 2012). The rise of the digital age and the consistent ability to access the internet has impacted these individuals by inundating them with stimuli. Although this instantaneous information available at their disposal has allowed for the formation of unique neural connections, it can lead people of this age group to be uninterested when experiences are under stimulating (Shams and Seitz, 2008; Starkey, 2012, Van Atteveldt et al., 2004).

One valuable consideration is the pedagogical styles used to help students learn and its connection to VR. Previous studies have linked constructivist approaches with VR being an addition to further enhance learning (Cheng and Wang, 2011; Huang et al., 2010; Sánchez et al., 2000). Constructivists approaches lean heavily on the ability for students to make connections with the material they’re learning and through their experiences, integrating this new knowledge/material into their ever-growing body of knowledge. Luckily in our present study our approaches are very constructivist in nature and lend themselves easily to VR use in higher education.

In our present study, we implemented VR experiences in a large introductory biology laboratory course at a public institution of higher education in the United States. We hypothesized that with VR experiences, students will not only be more engaged, but student performance on assessments will also be enhanced potentially through engaging multisensory learning. Our VR experiences used prepared materials freely available on the social media site YouTube (see materials section for additional information). We used 360-degree video, a form of virtual reality and many videos utilized were professionally produced (for example: Discovery and National Geographic). 360-degree video is footage shot with a special camera that immerses the viewer completely in a specific scene. The camera captures footage all around, below, above, from all angles. When the viewer is consuming the 360-degree video, they typically have a headset on and when the viewer’s head is pointed in a specific direction (leading the viewer to be able to look around in various directions for an immersive experience). Overall, through the implementation of VR experiences, we aim to enhance student attitudes, engagement, and performance for our students in foundational biological sciences material.

## Materials and Methods

### School Details and Demographics

The University of South Carolina (UofSC) is a state funded public four-year institution. The institution offers over 300 areas of study with majors, minors, and areas of emphasis covering a plethora of humanities, sciences (life and social), math, and art related fields, various certificate programs, Associate degrees, Bachelor of Arts, and Bachelor of Science degrees for numerous subjects for undergraduates. The school also has a thriving graduate program in many of the aforementioned subjects offering various degrees including master’s degrees and doctorate degrees. The UofSC system includes several professional schools such as medical schools (School of Medicine, Columbia SC; School of Medicine, Greenville SC), law school (UofSC School of Law), and a College of Pharmacy. The main campus is home to 35,296 total students, 26,753 of which are undergraduates. The largest major on campus is biological sciences which consists of 1,801 students. The course that was analyzed is not only a part of the biological sciences curriculum for the degree but also an optional part of the Carolina Core (general education) component. Demographics of the university can be found at the following link along with more details regarding the university as a whole: https://sc.edu/about/offices_and_divisions/institutional_research_assessment_and_analytics/documents/institutional_effectiveness/ipeds_data_feedback_reports/columbia/2021.pdf

### Class structure

Virtual reality (VR) was implemented in an introductory biology laboratory course (BIOL 102L) at the University of South Carolina. These labs typically meet once every week for two and a half hours and the maximum capacity of each lab is 24 students. Covering a wide variety of topics, the course is taken by students ranging from first year students to seniors (although most of our students are classified as first year). Students must either be currently enrolled or already completed the associated BIOL 102 lecture course to enroll in BIOL 102L. Student’s previous experience in biology varies widely, from high achieving advanced placement students to students that may not have taken any biology courses in their secondary school career. Each year we offer roughly 60 sections of BIOL 102L that are taught by the graduate students of the Department of Biological Sciences with a faculty member lab coordinator who trains the graduate student instructors, orders supplies, updates the curriculum, and handles all logistics of the implementation of the lab. For the semesters used in our study, Fall 2018 was comprised of 20 sections with a total of 471 students, Spring 2019 was comprised of 39 sections with a total of 900 students, Fall 2019 was comprised of 24 sections with a total of 540 students, and Spring 2020 was comprised of 39 sections with a total of 866 students.

### Class Curriculum and Format

BIOL 102L includes a wide variety of biological topics. Our major breaks up BIOL 101 and BIOL 102, the general principles of biological sciences, into two courses based on topics. BIOL 101 is typically based on cellular and molecular biology while BIOL 102 focuses on evolutionary, ecological, organismal, and basic anatomy and physiological biology. BIOL 102L is not directly linked to the course (the material covered is up to the instructor for BIOL 102 and with a wide variety of instructors), but generally covers the same topics. We also include topics such as experimental design and science communication to ensure our students have a firm foundation of how science operates. Our lab is delivered as a flipped course, meaning students are responsible for consuming and digesting material outside of the lab (and take a low-stakes quiz before lab starts). Typically, each lab session begins with a short formative in-lab assessment to test students on the previous lab’s material, followed by a short lecture and discussion by the graduate student instructor, and finally the students engage in the lab activities for the day (from designing and conducting various experiments to performing gallery walks of student work).

### VR Headsets

Each student in lab used a plastic VR headset in which they could insert their phone to view the VR experience (figure 1). We purchased and used the Onn Virtual Reality Smartphone headset. The headsets were purchased at a local store, transported, and distributed in a face-to-face physical lab on UofSC’s campus. A smartphone with the YouTube application installed was needed to be fully immersed in the VR experience. We also offered laptops for students that may not have a smartphone. While the experience wouldn’t be as immersive, they could still consume the material in a 2D format like any regular video. During our test semesters with VR (Fall 2019 and Spring 2020), out of 1,406 students there were zero students that didn’t have or use a smartphone for VR.

**Figure 1.**
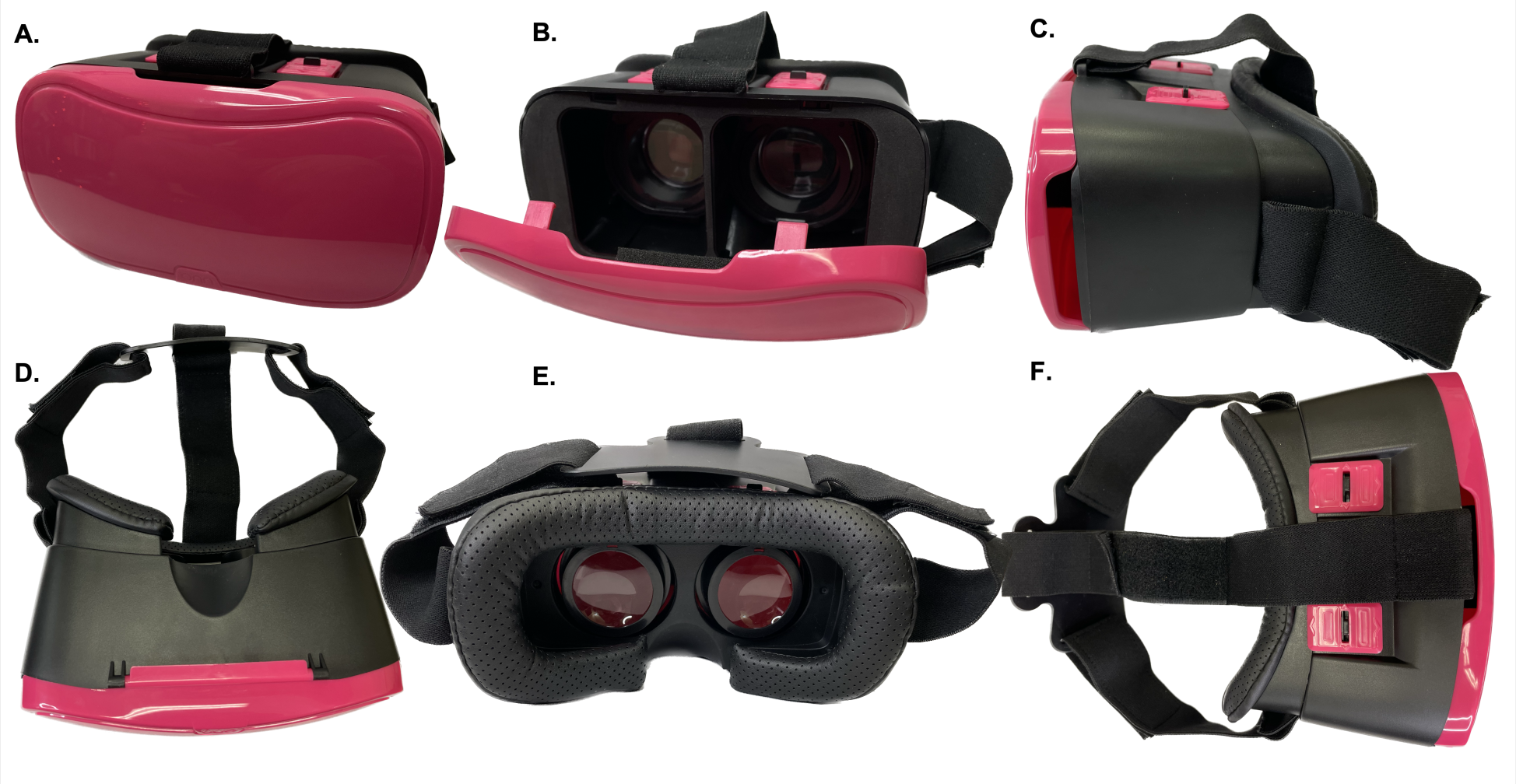
Image of a Virtual Reality Headset. To use the headset, one flips open the pink door of the headset, inserts a device that has the media displayed on the screen of the device, closes the door, and then places the headset on their head. The top of the headset has specific features that allow for the lenses to be adjusted closer or further from the viewer’s eyes and from side to side to allow for optimal viewing.

### VR Experiences

We selected specific 360-degree videos to be viewed by students in lab with the Onn VR headsets. These were pre-made, often extremely high quality, videos that were available free through YouTube. These videos were selected to supplement our curriculum and reinforce the topics that we were covering in lab that week. We selected videos focusing on topics that students had previously had issues engaging with, which include biodiversity, evolutionary biology, and plants. It’s important to note that each graduate student instructor was trained on how to use the Onn VR headsets and the amount of time that should be allotted in each lab session for the viewing of VR experiences. Each session would introduce the topic, and before beginning the lab activities of the day, students would spend roughly 15 minutes with the VR experience while engaging with students physically close to them in lab (so they aren’t just viewing the videos in isolation, but collaboratively with their peers). Below are the specific links to the 360-degree videos used in lab:

1. Surrounded by Wild Elephants in 4K 360: https://youtu.be/mlOiXMvMaZo
2. Wild With: Wolves (360 Video): https://youtu.be/vC8EVirfuMM
3. 360 Baby Pandas Nat Geo WILD: https://youtu.be/0XrH2WO1Mzs
4. Wild With: Bears (360 Video): https://youtu.be/lnh53i6RqQc
5. Lions 360 National Geographic: https://youtu.be/sPyAQQklc1s
6. Trip to the Caribbean: https://youtu.be/CIn-TjElcGw
7. Caribbean VR Experience Alpha: https://youtu.be/3DDqYdgvvxs
8. Galapagos 360: https://youtu.be/CcN6JkzM6Fk
9. 360 Galapagos Islands, Ecuador: https://youtu.be/gsPbaIeh_nQ
10. 360 Climbing Giants National Geographic: https://youtu.be/f7wTolIlK_s
11. 360 Video Tour of Congaree National Park: https://youtu.be/oUSioQ0Huu0
12. Krabi 360: Mangrove Forest: https://youtu.be/ZkNLqncV9xU

### Optional Survey

To assess student views and engagement with the VR experiences, we conducted an optional survey for each section. Using a standard Likert scale, we generated a Microsoft Form and encouraged students to fill out the survey once in the semester. As shown in the results section, we compiled these results to analyze the implementation of VR in BIOL 102L for semesters Fall 2019 and Spring 2020.

### Student Performance and Statistical Analysis

To analyze student performance in lab, we compared the grades of four different semesters: Fall 2018 and Spring 2019 that had no VR experience and Fall 2019 and Spring 2020 that contained VR experiences. Course grades along with withdrawal data were obtained through the publicly available data provided by the University of South Carolina Registrar’s office at the following link: https://sc.edu/about/offices_and_divisions/registrar/toolbox/grade_processing/grade_spreads/index.php.

We also analyzed each individual assessments that were linked directly with VR experiences to assess if VR enhanced student engagement. We performed Student T-Tests assuming equal variance to compare student performance on assessments and overall course grades between semesters with VR experience and those without.

Graphs and tables were constructed using R version 4.2.1 (“Funny -Looking Kid”) which is distributed under the terms of the GNU General Public License. R software is provided for free at www.R-project.org. R-studio was used to create the plots, along with ggplot2, and the ggprism theme (Dawson, 2022).

One plot type used to visualize the data of our study is the violin plot. This plot is useful for visualizing simple descriptive statistics (such as average and standard deviation) and includes a density plot to better see the distribution of the data. Violin plots can be read as follows: the wider the area of the geometric shape at a specific y axis value, the higher the probability of data being located at that specific value. For example, a plot that has wide geometric shapes at 70% and 80% y axis values has a higher probability of data being located at those values. We’ve included a boxpot in our violin plots to provide further distribution details (and a familiar plot). There’s also a point with error bars representing the mean along with standard error of the mean (the error bars).

## Results

### Students were enthusiastic about using virtual reality per an optional survey

The use of 360-degree video in BIOL 102L at UofSC was an optional but encouraged experience. At several points during the semester after completing a virtual reality experience, students were given the opportunity to complete a survey. A total of 679 students responded to the survey in Fall 2019 and Spring 2020 with a participation rate just under 50% (49.7%, n= 679). Students were provided several Likert scale statements and asked to select the response they most agreed with. Overall, students viewed the VR experience positively (and visualized in figure 2 below). We stated “I can better visualize some global biodiversity through this activity” to which 33.3% Strongly agreed (n=226), 47.6% Agreed (n=323), 13.5% Neutral (n=92), 2.9% Disagreed (n=20), and 2.7% Strongly disagreed (n=18). Our next statement was “This activity helped me engage with the topic” to which 39.2% Strongly agreed (n=266), 42.1% Agreed (n=286), 12.5% Neutral (n=85), 3.5% Disagreed (n=24), and 2.7% Strongly disagreed (n=18). We asked statements about the usage of VR in our lab and in other courses to assess if the technology is being utilized broadly. We stated “I would like to see more 360 video in lab” to which 36.7% Strongly agreed (n=249), 37.4% Agreed (n=254), 17.5% Neutral (n=119), 3.2% Disagreed (n=35), and 5.2% Strongly disagreed (n=22). Next, we wanted to assess whether the equipment/technology we were implementing caused students to have a negative experience. We stated “Technology distracted me from the topic at hand” to which 6% Strongly agreed (n=41), 9.9% Agreed (n=67), 14.5% Neutral (n=98), 44.6% Disagreed (n=303), and 25% Strongly disagreed (n=170). Our final statement was to determine the broad usage of VR in courses. We stated “I have used 360-degree videos/virtual reality in other courses” to which 5% Strongly agreed (n=34), 7.2% Agreed (n=49), 6.2% Neutral (n=42), 24.9% Disagreed (n=169), and 56.7% Strongly disagreed (n=385).

**Figure 2.**
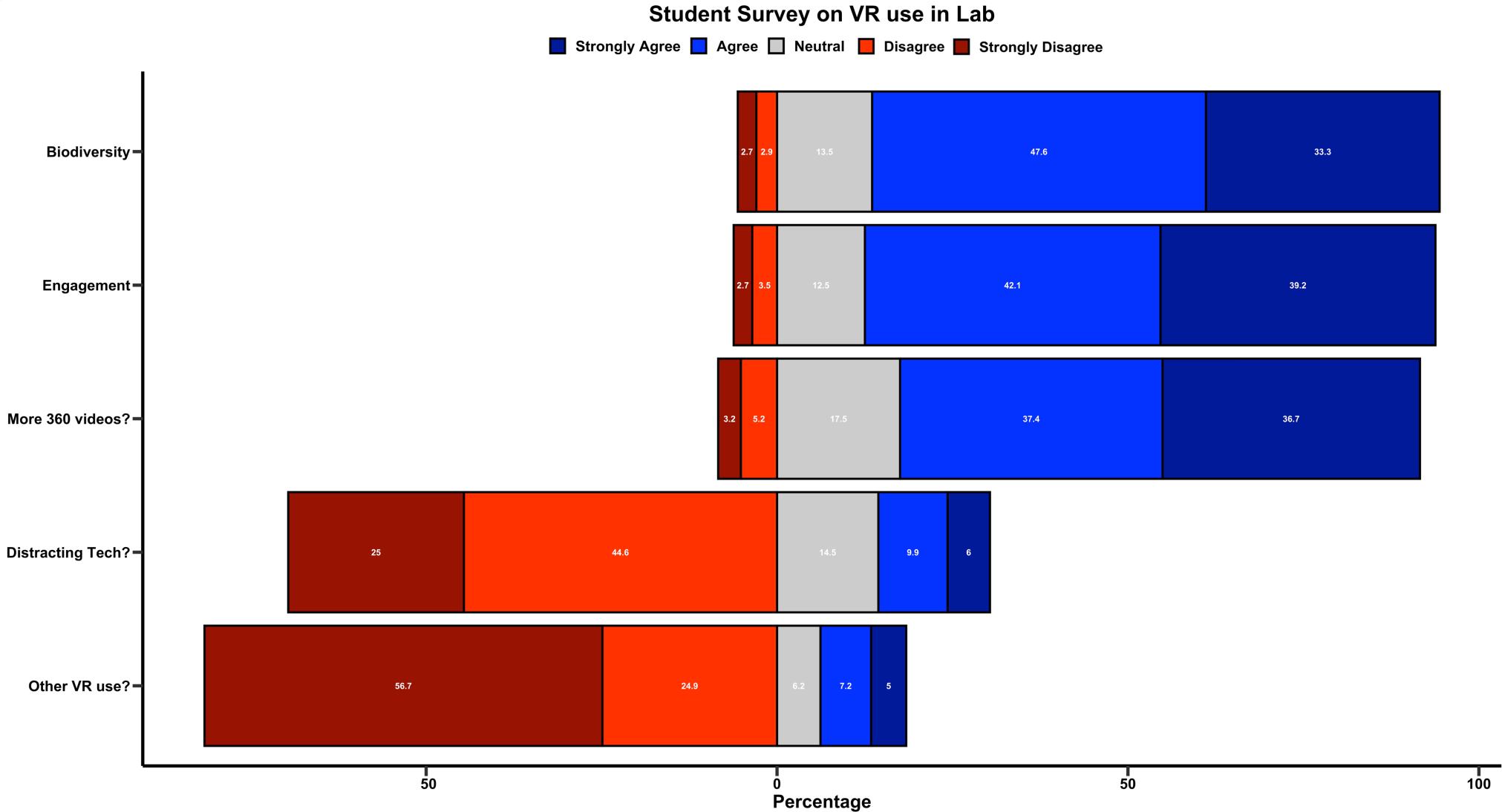
Survey data from students enrolled in BIOL 102L during the Fall 2019 and Spring 2020 semesters. A. Diverging bar graph visually representing the data. Each white number in the bar graph indicates the percentage of students that responded with that specific Likert scale choice. The optional survey was administered via Microsoft Forms directly after students had participated in a 360-degree virtual reality experience. Total number of participants was 679, with a participation rate of 49.7%. The results section includes the specific Likert statements (shortened here for appealing data visualization).

### Assessment performance and differing semesters

To assess the true impacts of virtual reality use, we analyzed several different regular assessments. Together, we analyzed formative quizzes (given at the beginning of lab to determine how students are doing on material from the previous lab), a summative lab practical (serving as a final exam in some respects), and we analyzed overall course grades. The formative quizzes were administered the lab section after the virtual reality experience took place and we only analyzed quizzes that were attempted (in other words, those students who did not complete the quiz were not included in the analysis). Specifically, we analyzed the biodiversity quiz, evolution quiz, and the plant quiz as they all had direct virtual reality experiences associated with the topic. The summative lab practical was selected to determine if virtual reality experience had impacts on retention of material throughout the course of the semester. Final course grades and DFW rates were also analyzed for semester with and without the use of virtual reality experiences. Fall and spring semesters were analyzed specifically (and distinctly) because there’s a traditional difference in course grades and performance overall. The major curriculum for biological sciences requires BIOL 101 (and BIOL 101L) and BIOL 102 (and BIOL 102L) but there’s no specific order in which they must be taken. Our advising model suggests that students who have not placed into calculus (via a placement assessment) should take BIOL 102 and BIOL 102L their first semester of school to keep pace with the major curriculum. Due to these suggestions, grades are typically significantly different from fall to spring semesters with spring semesters usually having higher course averages. These differences are also why we chose to analyze both fall and spring semesters with and without VR experiences.

### Virtual reality had minimal impact on biodiversity quiz performance

We first examined the quiz performance on the topic of diversity. Students had the opportunity to view several videos regarding biodiversity, mainly by showcasing specific exotic organisms (arguably one of most visually impactful aspects of biodiversity). Shown in figure 3, the quiz averages were very similar across all semesters examined: 87.69 for Fall 2018, 86.89 for Fall 2019 and 92.84 for Spring 2019, 89.55 for Spring 2020 (units: percentage). The overall distribution of the quiz grades had minimal changes between the 4 semesters. Note the general shape of figure 3 comes from the fact each quiz had low numbers of questions with no partial credit given (therefore, grade distributions were clumped at distinct locations).

**Figure 3.**
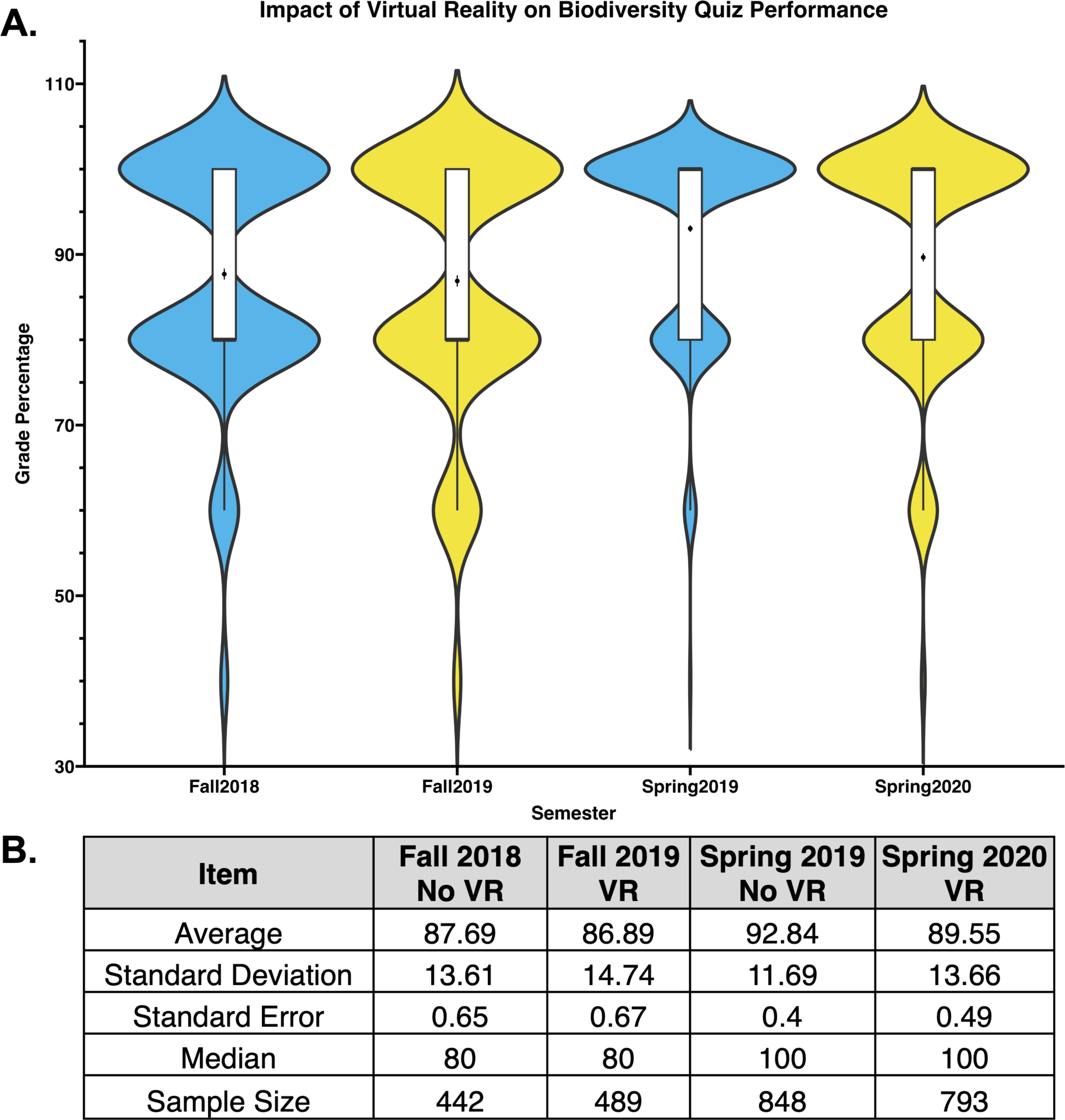
Biodiversity quiz performance was not impacted with use of virtual reality. All quiz grades from the biodiversity quiz were collected from Fall 2018, Spring 2019 with no VR experience and Fall 2019, Spring 2020 with VR experiences included in the course. A. Violin plot demonstrating the potential distribution of quiz grades for each semester. Black dot corresponds to the average quiz grade while the small black line represents the standard error of the mean. Boxplot included in each violin plot. In general, the areas of the plot with the largest width indicate relative density of grades at that specific percentage. B. Table summarizing the data from Figure 3A. Average, standard deviation, standard error, median, and the sample size are included. Blue color indicates a semester in which there was no VR experience while yellow color indicates a semester using VR.

### Virtual reality experiences were associated with enhanced evolution quiz performance for the spring semester

The formative quiz regarding evolution was analyzed to determine if VR experiences may influence performance. Visualized in figure 4, VR experiences in the fall semester had no significant impact on the quiz performance (fall 2018: 76.7%, fall 2019: 78.53%) while the VR experience was associated with an enhanced quiz performance in the spring semester (spring 2019: 77.86%, spring 2020: 84.58%). Statistical analysis of the grades using a Student’s T test found there was a statistically significant increase in grades from spring 2019 to spring 2020 (p-value: 1.05 × 10^-11^; t-stat: -6.75).

**Figure 4.**
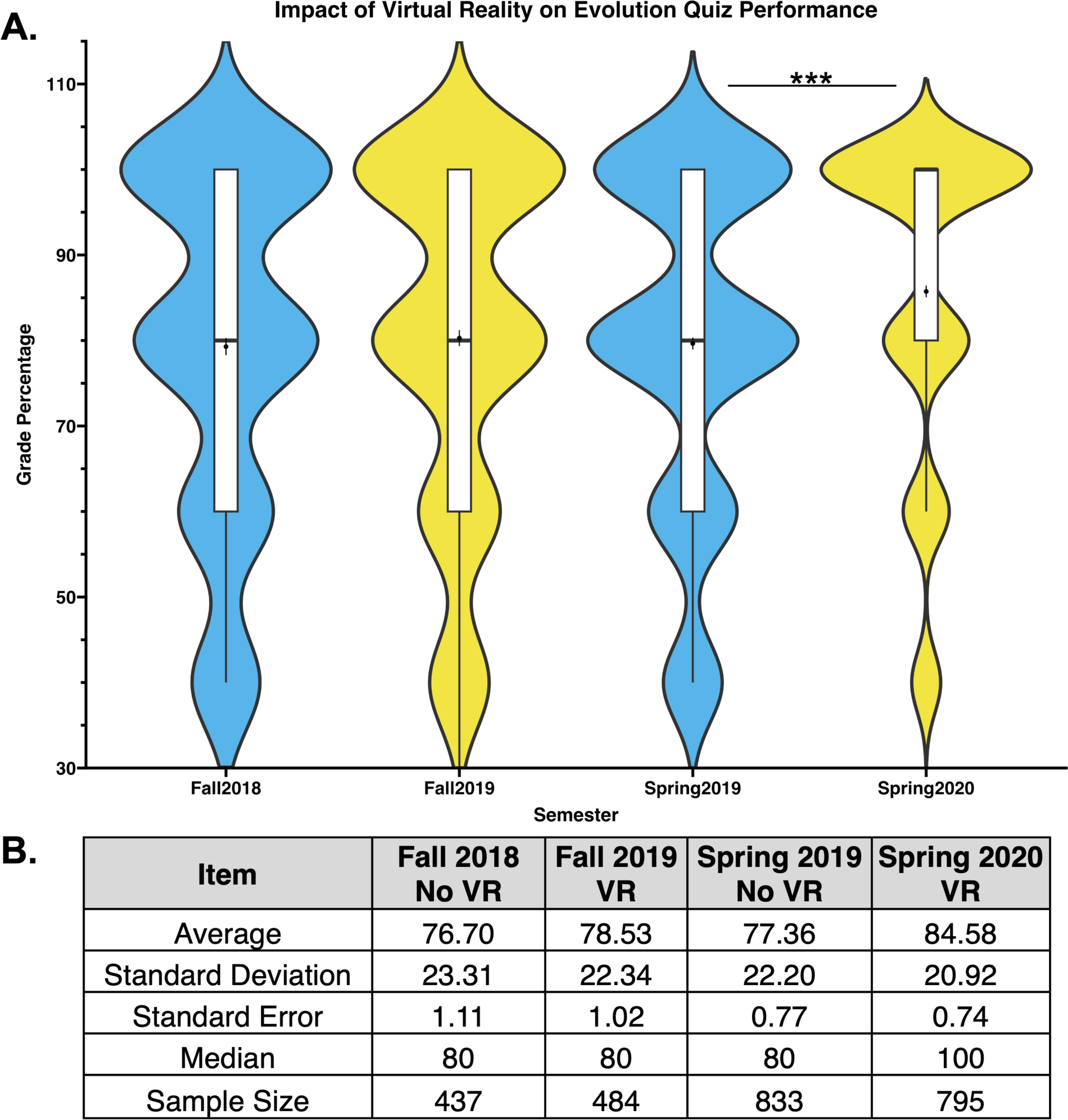
Evolution quiz performance improved in the spring semester with use of virtual reality. All quiz grades from the evolution specific quiz were collected from Fall 2018, Spring 2019 with no VR experience and Fall 2019, Spring 2020 with VR experiences included in the course. A. Violin plot demonstrating the potential distribution of quiz grades for each semester. Black dot corresponds to the average quiz grade while the small black line represents the standard error of the mean. Boxplot included in each violin plot. In general, the areas of the plot with the largest width indicate relative density of grades at that specific percentage. B. Table summarizing the data from Figure 4A. Average, standard deviation, standard error of the mean, median, and sample sizes are included. Blue color indicates a semester in which there was no VR experience while yellow color indicates a semester using VR.

### Virtual reality experiences were associated with improved plant quiz performance in each semester implemented

We also analyzed student performance on a formative quiz focused on the topic of plant biology. Traditionally, students have found the plant material of the course to be unengaging. Shown in figure 5, we analyzed the plant quiz performance in semesters with and without VR experiences. VR experiences were associated with a significant increase in quiz performance (averages for quiz: fall 2018: 86.18%, fall 2019: 92.03% and spring 2019: 87.81%, spring 2020: 94.59%). The distribution of the quiz grades was different between semesters with and without the VR experience, with an increase in high percentage grades (figure 5a). Statistical significance was determined by performing a Student’s T test and there was found to be a significant increase in grades from fall 2018 to fall 2019 and from spring 2019 to spring 2020 (fall data: p-value: 2.74 × 10^-10^, t-stat: -6.27; spring data: p-value: 1.09 × 10^-27^, t-stat: -11.04).

**Figure 5.**
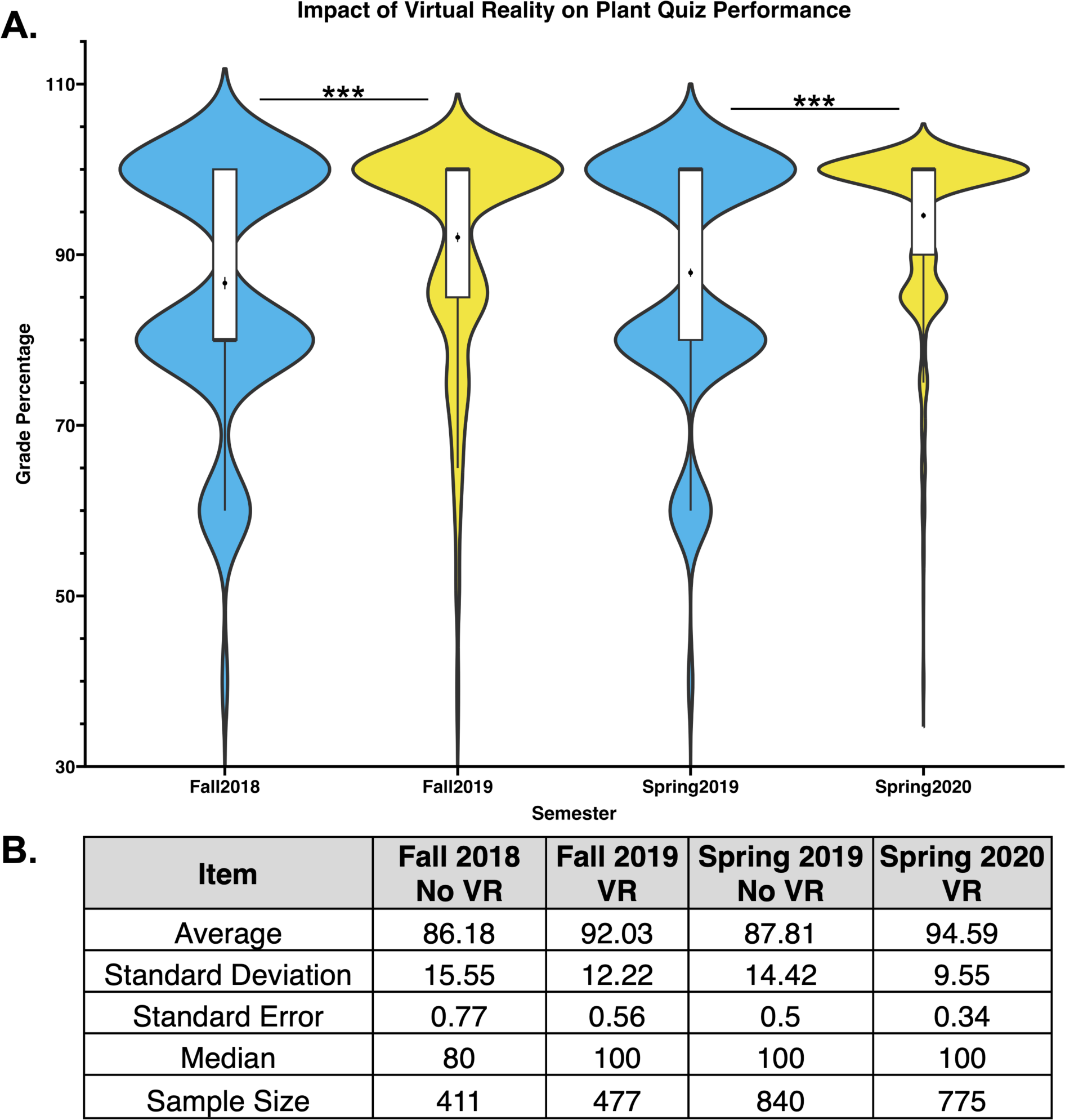
Plant quiz performance improved with use of virtual reality. All quiz grades from the plant quiz were collected from Fall 2018, Spring 2019 with no VR experience and Fall 2019, Spring 2020 with VR experiences included in the course. A. Violin plot demonstrating the potential distribution of quiz grades for each semester. Black dot corresponds to the average quiz grade while the small black line represents the standard error of the mean. Boxplot included in each violin plot. In general, the areas of the plot with the largest width indicate relative density of grades at that specific percentage. B. Table summarizing the data from Figure 5A. Average, standard deviation, standard error of the mean, median, and sample sizes are included. Blue color indicates a semester in which there was no VR experience while yellow color indicates a semester using VR.

### Summative assessment (lab practical) performance was enhanced in semesters with virtual reality experiences implemented

To analyze the impact of VR experiences on a summative assessment, we collected and analyzed performance on the lab practical. This summative assessment is administered at the end of each semester to determine collective knowledge gains. Shown in figure 6, VR experiences in lab were associated with improved assessment performance on the summative lab practical. For the fall semesters, the average grade on the practical was 75.48% for fall 2018 that had no VR experiences and increased to 79.44% for fall 2019 with VR experiences. Statistical significance was determined with a Student’s T test (p-value: 6.83 × 10^-7^, t-stat: -4.86). For spring semesters, the average grade on the practical was 78.53% for spring 2019 with no VR experience and 84.80% for spring 2020 that included VR experiences. Statistical significance was determined with a Student’s T test (p-value: 1.68 × 10^-34^, t-stat: -12.48). One situation to note is that in spring 2020, the lab was forced to move to an online modality due to the COVID-19 global pandemic. The lab practical was taken virtually that semester (along with the anatomy and physiology labs which contained no VR experiences).

**Figure 6.**
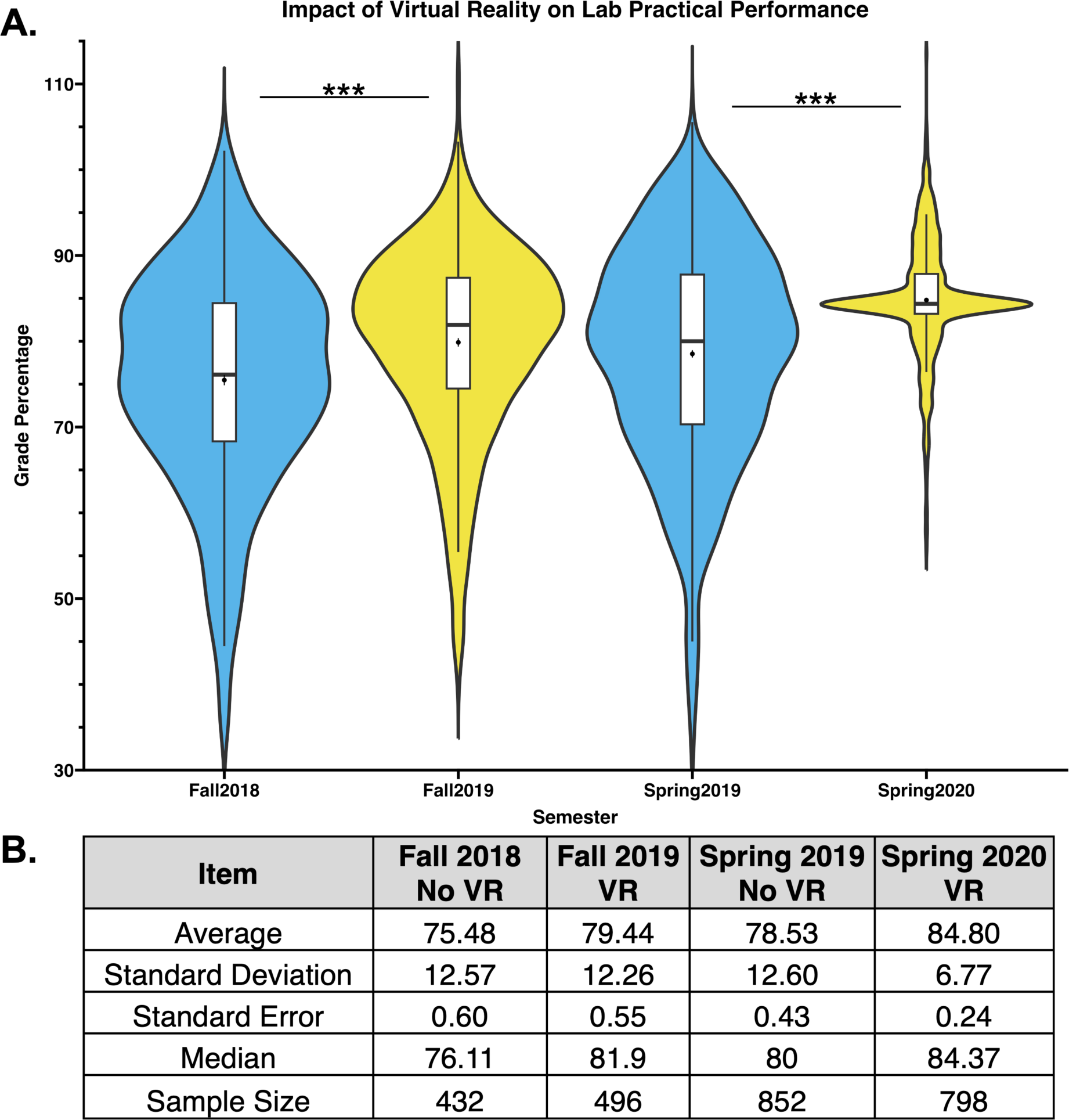
Lab practical performance improves with use of virtual reality. All lab practical grades were analyzed from Fall 2018, Spring 2019 with no VR experience and Fall 2019, Spring 2020 with VR experiences included in the course. A. Violin plot demonstrating the potential distribution of grades for each semester. Black dot corresponds to the average quiz grade while the small black line represents the standard error of the mean. Boxplot included in each violin plot. In general, the areas of the plot with the largest width indicate relative density of grades at that specific percentage. B. Table summarizing the data from Figure 6A. Average, standard deviation, standard error of the mean, median, and sample sizes are included. The lab practical is a summative assessment administered at the end of each academic semester to gauge retention of content. Blue color indicates a semester in which there was no VR experience while yellow color indicates a semester using VR.

### Virtual reality experiences were associated with higher final course grades

To assess overall impact of VR experiences on our course, we analyzed final course grades. Shown in figure 7, VR experiences in lab were associated with improved overall course grades for both semesters analyzed. For the fall semesters, the average course grade was 85.32% for fall 2018 that had no VR experiences and increased to 88.22% for fall 2019 with VR experiences. Statistical significance was determined with a Student’s T test (p-value: 6.92 × 10^-5^, t-stat: -3.82). For spring semesters, the average grade was 89.95% for spring 2019 with no VR experience and 91.77% for spring 2020 that included VR experiences. Statistical significance was determined with a Student’s T test (p-value: 8.53 × 10^-9^, t-stat: -5.67).

**Figure 7.**
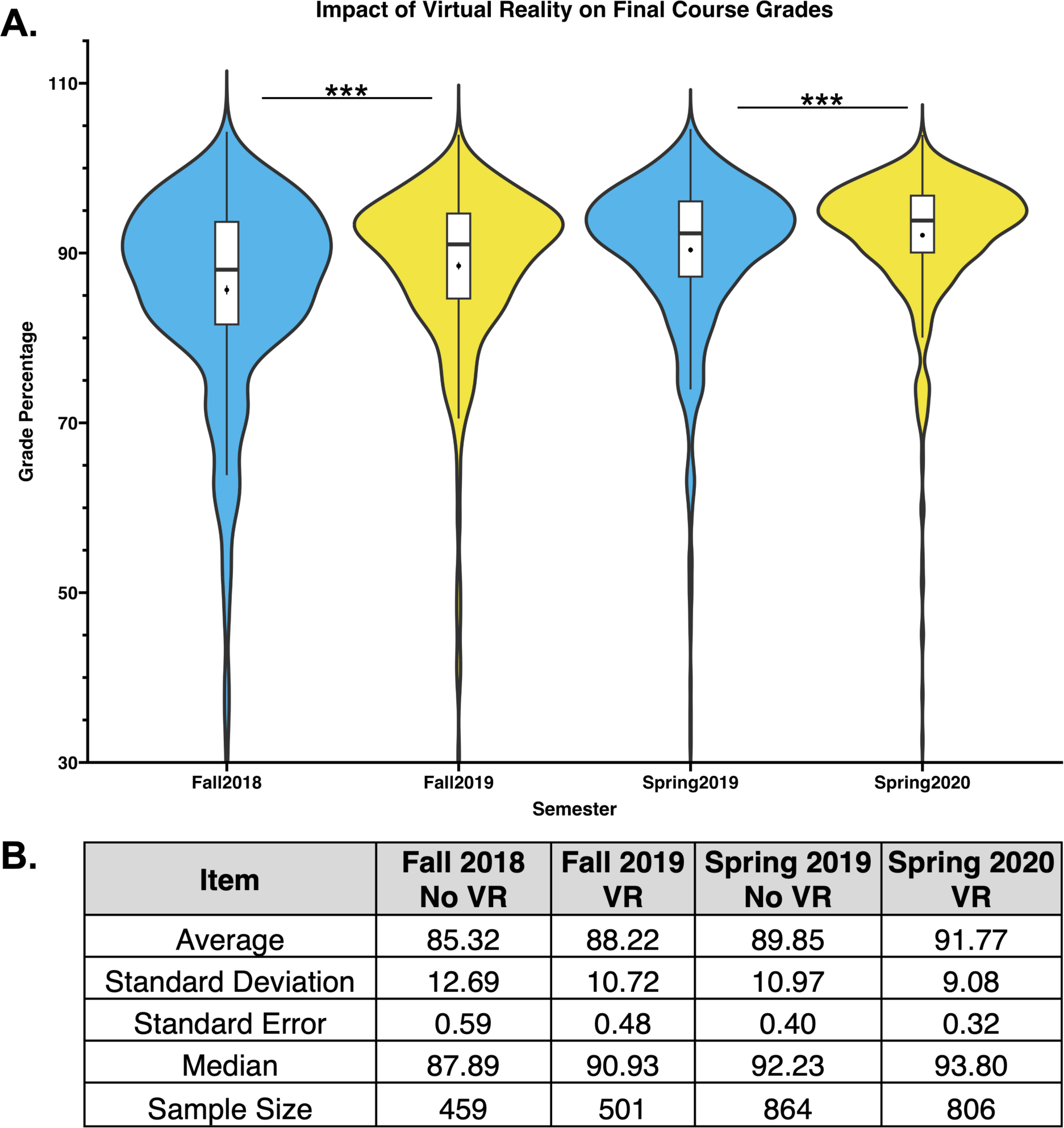
Final course grade performance improves with use of virtual reality. All grades were analyzed from Fall 2018, Spring 2019 with no VR experience and Fall 2019, Spring 2020 with VR experiences included in the course. A. Violin plot demonstrating the potential distribution of grades for each semester. Black dot corresponds to the average quiz grade while the small black line represents the standard error of the mean. Boxplot included in each violin plot. In general, the areas of the plot with the largest width indicate relative density of grades at that specific percentage. B. Table summarizing the data from Figure 7A. Average, standard deviation, standard error of the mean, median, and sample sizes are included. Blue color indicates a semester in which there was no VR experience while yellow color indicates a semester using VR.

### Semesters with virtual reality experiences exhibited reduced rates of D and F rates

Demonstrated in figure 8, VR use was associated with greatly decreased rates of D and F with minimal impact on withdrawal rates in the fall semester, and a potential reduction for the spring semester (although midway through spring 2020 the pandemic forced face to face education to move online, potentially impacting withdrawal rates). For fall 2018 the D rate was 4.03% with no VR experience and 0.56% for fall 2019 with VR experiences, a decrease of 86.1%. For spring 2019 the D rate was 1.11% with no VR experience and 0.11% for spring 2020 with VR experiences, a decrease of 90%. For fall 2018 the F rate was 4.46% with no VR experience and 2.41% for fall 2019 with VR experiences, a decrease of 45.96%. For spring 2019 the F rate was 2.33% with no VR experience and 0.81% for spring 2020 with VR experiences, a decrease of 65.23%. The withdrawal rate for fall 2018 was 2.97% with no VR experience and 3.14% for fall 2019 with VR experiences. The withdrawal rate for spring 2018 was 2.89% with no VR experience and 1.39% for fall 2019 with VR experiences.

**Figure 8.**
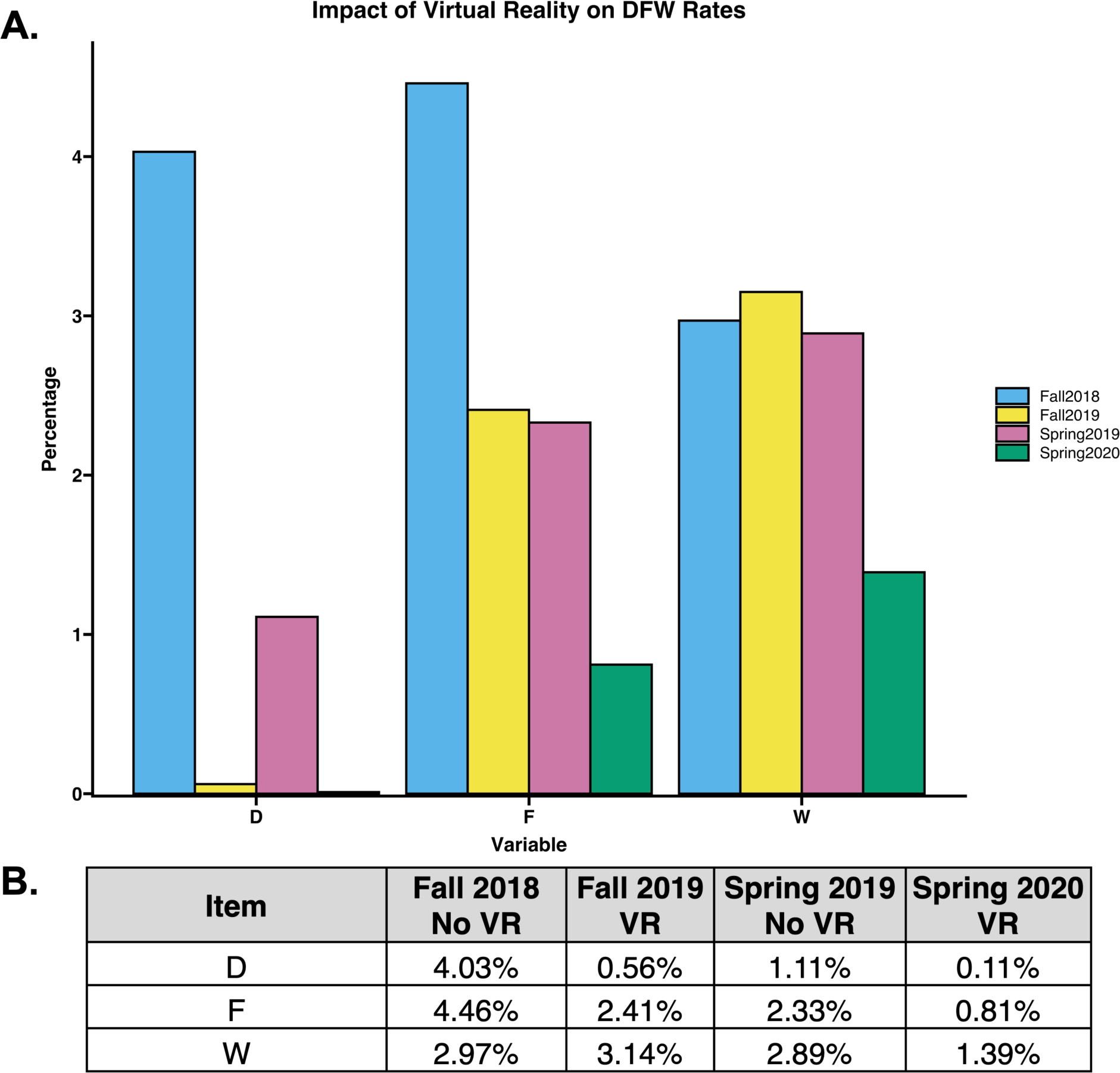
DF rates improve with use of virtual reality, W rates improved in spring semester with use of virtual reality. D, F, and W rates were analyzed from Fall 2018, Spring 2019 with no VR experience and Fall 2019, Spring 2020 with VR experiences included in the course. A. A bar graph for each category (D, F, and W) is presented. B. Table summarizing the data from Figure 8A.

## Discussion

In our current study, we’ve assessed the impact of VR experiences on various formative assessments, a summative assessment, DFW rates, and overall course performance. Collected over 4 semesters and over a hundred sections of lab, our data suggests VR experiences in an introductory lab course is associated with enhanced student engagement, student attitudes, and overall course performance.

Analyzing the data from our survey, students had a positive experience with VR in lab. From enhancing engagement with specific topics to better visualizing global biodiversity, students seemed to agree that the experience was beneficial (Figure 2). We even assessed their views on putting more VR experiences in lab and students were enthusiastically in favor, 74.1% of students either strongly agreeing or agreeing (Figure 2). Previous studies have shown that technology use and potential motion sickness may occur and result in undesirable experiences for students/users of VR in general (Saradakis et al., 2020). To help address the technology aspects, we asked students if technology distracted them from the topic and 69.6% of students either disagreed or strongly disagreed (Figure 2). While students were enthusiastic about VR experiences in lab, our survey results suggest it’s not something they’ve encountered in many other courses (81.6% either disagreed or strongly disagreed to the question “I have used 360-degree videos/virtual reality in other courses”). Overall, the reception of the VR experience by students was extremely positive and came with praise from the instructor, which may have helped students engage fully (engagement helps students meet their academic goals). (Hourigan, 2013; McClenney, 2007; Persell et al., 2008).

Students saw the VR experiences positively and we next analyzed student performance on various assessments to determine if there were any associations with enhanced performance and VR usage. A few studies have been published that indicate student performance was enhanced with the implementation of virtual reality experiences in higher education, although none looking specifically at large numbers over multiple years in a biological sciences curriculum and/or included formative and summative assessment performance (Bennett and Saunders, 2019; Lee and Wong, 2014; Makransky and Lilleholt, 2018; Shim et al., 2003; Zhao et al., 2020). The first VR experience used in the course was one focused on global biodiversity (Figure 3). Interestingly, there was no associated impact on biodiversity quiz performance with VR usage. We speculate that this result may be simply because it was the first VR experience so students may have spent more time trying to make the equipment function instead of focusing on the content. Alternatively, the 360-degree videos we selected were about large fauna found globally (and were relatively popular organisms, such as wolves, pandas, and lions). Due to this popularity, students may already have a sense of large organism biodiversity and thus why the VR experience wasn’t associated with any specific enhancement in quiz performance (Figure 3).

The next assessment focused on the theory of evolution and Hardy-Weinberg equilibrium (a concept that challenges students). We found that the fall semesters experienced no impact on quiz performance with the implementation of VR while the spring semester had a statistically significant increase in quiz grades associated with the use of VR (Figure 4). The VR experience used focuses on places of historic importance; for example, we show the students what life is like on the Galápagos Islands, the place credited for the inception of Darwin’s famous theory of evolution. While the setting may have been important for spring semester students, those in the fall may have needed additional information for the VR experience to potentially make a difference in performance. Another important consideration is the topic of Hardy-Weinberg equilibrium. Traditionally the problems assigned need a solid foundation of math, and our fall students may not have that solid foundation (thus potentially resulting in no significance with VR experiences).

Another subject students traditionally struggle with is plant biology. We found a statistically significant increase in plant quiz performance associated with VR implementation of both fall and spring semesters analyzed (Figure 5). These experiences showcased the amazing biodiversity of plants and helped students see the importance of plants.

To analyze the potential impacts of VR on a summative assessment, we analyzed performance on the lab practical (Figure 6). Again, we found a statistically significant increase in performance for both fall and spring semesters. We analyzed the overall associated potential impacts of VR usage on overall course grades and found a statistically significant increase in semesters that implemented a VR experience.

Then to analyze potential impacts on final course grades, we compared semesters with and without VR usage (Figure 7). Here we uncovered that both fall and spring semesters had a statistically significant increase in student grades in semesters that had a VR experience than in semesters without a VR experience. While several factors could potentially influence final course grade, including admission standard changes or advisement changes, we’re optimistic that VR experiences had a positive impact on student course grades and engagement as has been found in other studies using VR in education (Lee and Wong, 2014; Makransky and Lilleholt, 2018; Shim et al., 2003; Zhao et al., 2020)

Lastly, we assessed the D, F, and withdrawal rates (Figure 8). Both D and F rates had significantly lower rates in semesters that implemented VR experiences. Fall semester withdrawal rates weren’t significantly altered while there was an associated decrease in withdrawal rates for the spring semester. While these results are encouraging, it should be noted that specifically for withdrawal rates, the spring semester did experience a shutdown due to the global pandemic and a shift to virtual learning.

Overall, our results demonstrate a potential significant impact of virtual reality experiences in an introductory college course. Our findings support our hypothesis that VR experiences will enhance student engagement, attitudes, and performance. While two formative assessments: the biodiversity quiz and fall semester evolution quiz, didn’t have enhanced performance with VR, we speculate a more refined experience could potentially lead to enhanced performance for these quizzes as well. Virtual reality has already been shown in various studied to enhance student engagement (Huang and Liaw, 2018; Hu-Au and Lee, 2017); in our present study, we link student performance on assessment in a large sample size over multiple years of study to add even further value to the implementation of virtual reality in higher education. Once used primarily as a training technique for pilots and associated with next level gaming, virtual reality has a place in higher education where it can help cement concepts for our students. Multi sensory processing and the four stages of learning are all enhanced through the engagement of sight and sound with the implementation of VR in the classroom (Quak et al., 2015; Shams and Seitz, 2008). In future work, we’ll investigate potential uses of augmented reality (potentially with the use of the popular social media app Snapchat) and even the gamification of these virtual reality experiences in an active, social learning environment. While VR experiences may not be for all students (i.e. motion sickness, Saradakis et al., 2020), it can serve extremely important roles in supplementing our curriculum in ways to further enhance engagement and performance.

